# Motif-centric phosphoproteomics to target kinase-mediated signaling pathways

**DOI:** 10.1101/2021.07.02.450911

**Authors:** Chia-Feng Tsai, Kosuke Ogata, Naoyuki Sugiyama, Yasushi Ishihama

**Affiliations:** Graduate School of Pharmaceutical Sciences, Kyoto University, Kyoto 606-8501, Japan; Laboratory of Clinical and Analytical Chemistry, National Institute of Biomedical Innovation, Health and Nutrition, Ibaraki, Osaka, 567-0085, Japan

**Keywords:** Motif-centric, Phosphoproteome, Kinase, IMAC

## Abstract

Identifying cellular phosphorylation pathways based on kinase-substrate relationships is a critical step to understanding the regulation of physiological functions in cells. Mass spectrometry-based phosphoproteomics workflows have made it possible to comprehensively collect information on individual phosphorylation sites in a variety of samples. However, there is still no generic approach to uncover phosphorylation networks based on kinase-substrate relationships in rare cell populations. Here, we describe a motif-centric phosphoproteomics approach combined with multiplexed isobaric labeling, in which *in vitro* kinase reaction is used to generate the targeted phosphopeptides, which are spiked into one of the isobaric channels to increase detectability. Proof-of-concept experiments demonstrate selective and comprehensive quantification of targeted phosphopeptides by using multiple kinases for motif-centric channels. Over 7,000 tyrosine phosphorylation sites were quantified from several tens of µg of starting materials. This approach enables the quantification of multiple phosphorylation pathways under physiological or pathological regulation in a motif-centric manner.

**Motivation:** Sensitivity for detecting phosphopeptides with a particular phosphorylation motif is limited, especially for tyrosine phosphopeptides.

## Introduction

Protein kinase-mediated phosphorylation on serine, threonine and tyrosine residues is one of the most ubiquitous post-translational modifications (PTMs). Signaling cascades via protein phosphorylation play key roles in multiple cellular processes in mammals, including intra- and intercellular signaling, protein synthesis, gene expression, cell survival and apoptosis (Cohen, 2002; Hunter, 2000). The relative abundances of phosphoserine (pS), phosphothreonine (pT), and phosphotyrosine (pY) sites in the human proteome have been estimated to be 90:10:0.05 based on the traditional method of ^32^P labeling (Hunter and Sefton, 1980). There are many possible reasons for the extreme paucity of pY sites compared to pS and pT sites in mammals, including the fact that tyrosine kinases are activated only under certain conditions, and that the high activity of protein tyrosine phosphatase (PTP) leads to a short half-life of pY sites (Hunter, 2009).

Great advances in the analytical workflows of shotgun phosphoproteomics, in which metal affinity chromatography is integrated with LC/MS/MS, have made it possible to identify more than 30,000 phosphorylation sites (Bekker-Jensen et al., 2017; Hogrebe et al., 2018; Mertins et al., 2018). In general, LC/MS/MS has an identification bias toward the more abundant phosphopeptides in a sample, whereas in kinase substrates, sequence features such as Pro-directed, basophilic, acidophilic and tyrosine-containing motifs have an important influence (Villen et al., 2007). Also, since biological importance does not necessarily correlate with protein expression levels, it is possible that important signals are transduced via specific kinases that are expressed at extremely low levels. Therefore, advanced pre-fractionation or enrichment methods prior to LC/MS/MS are needed to identify a wide range of kinase substrates (Tsai et al., 2014). Especially for low-abundance pY peptides, MS detectability is affected by the ionization suppression caused by the presence of more abundant pS and pT peptides in the complex phosphoproteomes. However, the combination of metal-affinity chromatography with immunoaffinity purification using pY antibody (Abe et al., 2017) or a recently developed SH2 domain-derived pY-superbinder (Bian et al., 2016; Dong et al., 2017) has been reported to increase the identification number of pY peptides. In addition, immunoaffinity-based methods using multiple antibodies have been developed for identification and quantitation of phosphopeptides derived from proteins in various pathways or pY peptides derived from tyrosine kinases (Stokes et al., 2012). Nevertheless, a large amount of starting material (mg to ten mg level) is generally necessary for deep tyrosine phosphoproteome analysis.

One of the major advantages of multiplexed isobaric tandem mass tag (TMT)-based methods for relative quantitation is that the differentially labeled peptides appear as a single peak at the MS1 level (Thompson et al., 2003), enhancing the detectability of low-abundance peptides. TMT strategies using a large amount of relevant “boosting” (or “carrier”) peptides labeled with one or several TMT channels have been successfully used for single-cell proteomics analysis (Budnik et al., 2018; Dou et al., 2019; Tsai et al., 2020). For instance, Lian *et al*. developed a BASIL (Boosting to Amplify Signal with Isobaric Labeling) strategy to quantify >20,000 phosphorylation sites in human pancreatic islets. However, the identification number of pY peptides was less than 1%. Recently, Chua *et al*. developed a BOOST (Broad-spectrum Optimization Of Selective Triggering) method in which pervanadate (tyrosine phosphatase inhibitor)-treated cells were used as a boosting channel to increase the detectability of pY peptides (Chua et al., 2020). The BOOST method coupled with antibody-based pY enrichment could quantify over 2300 unique pY peptides. However, the required amount of starting material was in the mg range, making it difficult to apply to small samples such as clinical specimens, which often contain less than 100 μg of extractable material.

We previously developed an LC/MS/MS-based *in vitro* kinase assay using dephosphorylated lysate proteins as the substrate source for *in vitro* kinase reaction to profile human protein kinomes (Imamura et al., 2014). A total of 175,574 direct kinase substrates were identified from 354 wild-type protein kinases, 21 mutant protein kinases, and 10 lipid kinases (Sugiyama et al., 2019). In addition, we utilized the *in vitro* kinase reactions with CK2, MAPK and EGFR to generate phosphopeptides with targeted motifs to measure the phosphorylation stoichiometry of >1,000 phosphorylation sites, including 366 low-abundance tyrosine phosphorylation sites (Tsai et al., 2015).

In the present study, we aimed to develop a motif-centric TMT approach in which phosphopeptides having targeted sequence motifs are generated by *in vitro* kinase reaction for the boosting TMT channel in order to increase the detectability of kinase substrates, including tyrosine kinase substrates, without immunoaffinity enrichment. To demonstrate the feasibility of this strategy, phosphopeptides with targeted motifs of CK2, PKA, CDK1, ERK2, JNK1, p38a, SRC and EGFR were used for the boosting TMT channel to monitor the perturbation of kinases-mediated signaling pathways by tyrosine kinase inhibitor treatment.

## STAR★Methods

| REAGENT or RESOURCE | SOURCE | IDENTIFIER |
| --- | --- | --- |
| **Biological Samples** |  |  |
| HeLa S3 | ATCC | Catalog: CCL-2.2 |
| **Critical Commercial Reagents and Assays** |  |  |
| Triethylammonium bicarbonate | Sigma | Catalog: T7408 |
| Phosphatase Inhibitor Cocktail 2 | Sigma | Catalog: P5726 |
| Phosphatase Inhibitor Cocktail 3 | Sigma | Catalog: P0044 |
| BCA Protein Assay Kit | Thermo Scientific<br>Pierce | Catalog: 23225 |
| Sodium deoxycholate | FUJIFILM Wako | Catalog: 190-08313 |
| Sodium lauroyl sarcosinate | FUJIFILM Wako | Catalog: 198-14745 |
| Iron-(III) chloride | FUJIFILM Wako | Catalog: 091-00872 |
| Dithiothreitol | Thermo Scientific | Catalog: 20291 |
| Iodoacetamide | Thermo Scientific | Catalog: A3221 |
| Lysyl endopeptidase | FUJIFILM Wako | Catalog: 129-02541 |
| Sequencing-grade modified trypsin | Promega | Catalog: V517 |
| Ni-NTA silica resins | QIAGEN | Catalog: 31314 |
| Empore SDB-XC membrane disks | CDS | Catalog: 13-110-020 |
| CK2α1/$\beta$ (CSNK2A1/B) | Carna Biosciences | Catalog:05-184 |
| CK2α2/$\beta$ (CSNK2A2/B) | Carna Biosciences | Catalog:05-185 |
| PKACα(PRKACA) | Carna Biosciences | Catalog:01-127 |
| ERK2 (MAPK1) | Carna Biosciences | Catalog:04-143 |
| EGFR (ERBB1) | Carna Biosciences | Catalog:08-115 |
| SRC | Carna Biosciences | Catalog:08-173 |
| JNK1(MAPK8) | Carna Biosciences | Catalog:04-163 |
| CDK1 (CDC2/CycB1) | Carna Biosciences | Catalog:04-102 |
| p38α(MAPK14) | Carna Biosciences | Catalog:04-152 |
| TMTsixplex™ | Thermo Scientific | Catalog:90061 |
| **Software and Algorithms** |  |  |
| Maxquant | PMID: 27809316 | https://www.maxquant.org/ |
| Perseus | PMID: 27348712 | https://maxquant.net/perseus/ |
| **Deposited Data** |  |  |
| jPOST Repository | PMID: 30295851 | https://repository.jpostdb.org |

### Tryptic peptides from HeLa cell lysate

HeLa S3 cells were cultured in DMEM containing 10% FBS and 100 µg/mL kanamycin. For isobaric acidophilic motif-centric phosphoproteomes, cells were not stimulated (mock) or were stimulated with 10 µM CK2 inhibitor (CX-4945) for 30 mins. For isobaric basophilic motif-centric phosphoproteomes, cells were not stimulated (mock) or were stimulated with 10 µM PKA activator (forskolin) for 30 mins. For isobaric tyrosine and multiple motif-centric phosphoproteome, cells were treated with 10 µM EGF, 10 µM EGF/10 µM afatinib and 500 µM pervanadate (pH 10 with 0.14% H_2_O_2_), respectively, for 30 mins before harvesting. Two biological replicates were performed. Cells were washed three times with ice-cold phosphate-buffered saline (PBS, 0.01 M sodium phosphate, 0.14 M NaCl, pH 7.4) and then lysed in lysis buffer containing 12 mM sodium deoxycholate (SDC), 12 mM sodium lauroyl sarcosinate (SLS) in 100 mM triethylammonium bicarbonate (TEABC). Protein concentration was determined by means of BCA protein assay. The lysates were digested based on the reported phase-transfer surfactants (PTS) protocol (Masuda et al., 2008). The digested peptides were desalted on SDB-XC StageTips (Rappsilber et al., 2007).

### H3In vitro kinase reaction

For acidophilic, Pro-directed and tyrosine kinase reactions, the tryptic peptides were dissolved in 40 mM Tris-HCl (pH 7.5) and incubated with each kinase (0.2 µg CK2, ERK2, JNK1, p38α, CDK1 or SRC) at 37 °C for *in vitro* kinase reaction in the presence of 1 mM ATP and 20 mM MgCl_2_. For EGFR kinase reaction, tryptic peptides were firstly passed through SCX StageTips (Rappsilber et al., 2007) to remove afatinib. Eluted peptides were further desalted on SDB-XC StageTips. Then, the desalted peptides were dissolved in 40 mM Tris-HCl (pH 7.5) and incubated with EGFR (0.2 µg) for *in vitro* kinase reaction in the presence of 1 mM ATP and 4 mM MnCl_2_ at 37 °C. For basophilic kinases such as PKA, the lysates were loaded onto a 10 kDa ultrafiltration device (Amicon Ultra, Millipore). The device was centrifuged at 14,000 g to remove the detergents. Subsequently, the original lysis buffer was replaced with 40 mM Tris-HCl (pH 7.5) followed by centrifugation. Then, the proteins were incubated with 0.2 µg PKA for *in vitro* kinase reaction in the presence of 1 mM ATP and 20 mM MgCl_2_. After the kinase reaction, the proteins were reduced with 10 mM DTT for 30 mins at 37 °C and alkylated with 50 mM iodoacetamide in the dark for 30 mins at 37 °C. The resulting samples were digested by Lys-C (1:100, w/w) at 37 °C for 3 hr followed by trypsin (1:50, w/w) overnight at 37 °C. All the peptides were desalted on SDB-XC StageTips.

### TMT labeling for purified phosphopeptides

The desalted peptides were dissolved in 200 mM HEPES (pH 8.5). Then, the resuspended phosphopeptides were mixed with TMT reagent dissolved in 100% ACN for 1 hr. The labeling reaction was stopped by adding 5% hydroxylamine for 15 minutes, followed by acidification with TFA. All the phosphopeptides labeled with each multiplexed TMT reagent were mixed into the same tube and the mixture was diluted to reduce the concentration of ACN to below 5%. The TMT-labeled peptides were desalted on SDB-XC StageTips.

### Immobilized metal ion affinity chromatography (IMAC)

The procedure for phosphopeptides purification with an Fe^3+^-IMAC tip was as described previously (Tsai et al., 2014; Tsai et al., 2015) with minor modifications. In brief, buffer consisting of 50 mM EDTA in 1 M NaCl was used for removing Ni^2+^ ions. Then, the metal-free NTA was activated by loading 100 mM FeCl_3_ into the IMAC tip. The Fe^3+^-IMAC tip was equilibrated with 1% (v/v) acetic acid at pH 3.0 prior to sample loading. Tryptic peptides from HeLa lysates were reconstituted in 1% (v/v) acetic acid and loaded onto the IMAC tip. After successive washing steps with 1% (v/v) TFA in 80% ACN and 1% (v/v) acetic acid, the IMAC tip was inserted into an activated SDB-XC StageTip. Then, the bound phosphopeptides were eluted into the SDB-XC StageTip by adding 200 mM NH_4_H_2_PO_4_. The eluted phosphopeptides were desalted on the SDB-XC StageTip.

### LC/MS/MS analysis

NanoLC/MS/MS analyses were performed on an Orbitrap Fusion Lumos Tribrid mass spectrometer (Thermo Scientific, San Jose, CA), which was connected to a Thermo Ultimate 3000 RSLCnano system (Germering, Germany) and an HTC-PAL autosampler (CTC Analytics, Zwingen, Switzerland). Peptide mixtures were loaded onto and separated on self-pulled needle columns (150 mm length × 100 μm ID) packed with Reprosil-Pur 120 C18-AQ material (3 μm, Dr. Maisch GmbH, Amerbuch, Germany) or a 2-m long C18 monolithic silica capillary column (Iwasaki et al., 2010). The mobile phases consisted of (A) 0.5% acetic acid and (B) 0.5% acetic acid and 80% acetonitrile. Peptides were separated through a gradient from 17.5 % to 45% buffer B at a flow rate of 500 nL/min. Full-scan spectra were acquired at a target value of 4×10^5^ with a resolution of 60,000. Data were acquired in a data-dependent acquisition mode using the top-speed method (3 sec). The peptides were isolated using a quadrupole system (the isolation window was 0.7). The MS^2^ analysis was performed in the ion trap using CID fragmentation with a collision energy of 35 at a target value of 1 × 10^4^ with 100 mS maximum injection time. The MS^3^ analysis was performed for each MS^2^ scan acquired by using multiple MS^2^ fragment ions isolated by an ion trap as precursors for the MS^3^ analysis with a multinotch isolation waveform(McAlister et al., 2014). HCD fragmentation was used for MS^3^ scan with an NCE of 65%, and the fragment ions were detected by the Orbitrap (resolution 15000). The AGC target was 5×10^4^ with a maximum ion injection time of 22 ms. The raw data sets have been deposited at the ProteomeXchange Consortium (http://proteomecentral.proteomexchange.org) via the jPOST partner repository (http://jpostdb.org) (Moriya et al., 2019) with the dataset identifier JPST001027 (PXD026996).

### Database search

The raw MS/MS data were processed with MaxQuant (Cox and Mann, 2008; Tyanova et al., 2016a). Peptide search with full tryptic digestion and a maximum of two missed cleavages was performed against the SwissProt human database (20,102 entries). The mass tolerance for precursor and MS^3^ ions was 4.5 ppm, whereas the tolerance for MS^2^ ions was 0.5 Th. Acetylation (protein *N*-terminal), oxidation (M) and phospho (STY) were set as variable modifications and carbamidomethyl (C) was set as a fixed modification. The quantitation function of reporter ion MS^3^ (6-plexed TMT) was turned on. The false-discovery rate (FDR) was set to 1% at the level of PSMs and proteins. A score cut-off of 40 was used for identified modified peptides. The abundances of TMT were log_2_-transformed and further analyzed by Perseus (Tyanova et al., 2016b) for statistical evaluation. The PSP logo generator (Hornbeck et al., 2015) was used for sequence motif analysis.

## Results

### Workflow for isobaric motif-centric phosphoproteome analysis

We previously developed a motif-centric approach (Tsai et al., 2015) in which dephosphorylation and isotope tagging are integrated with *in vitro* kinase reaction to improve the sensitivity and reproducibility for determining the absolute phosphorylation stoichiometry of targeted kinase substrates. However, the number of phosphosites commonly identified in endogenous and motif-centric phosphopeptides was not as large as expected, because some endogenous signals with specific kinase motifs are below the detection limit. In this study, we employed not isotopic but isobaric tagging to develop a motif-centric approach in which the same peptides from different samples were labeled with multiplexed TMT reagents and assembled as a single peak at the MS^1^ level to increase the sensitivity. In addition, we set one of the TMT channels for signal boosting, utilizing phosphopeptides having targeted sequence motifs generated by *in vitro* kinase reaction to increase the detectability of targeted kinase substrates. The entire workflow is shown in Figure 1. The motif-centric peptides are generated by *in vitro* kinase reactions using Pro-directed, acidophilic, basophilic or tyrosine kinase (Figure 1a). The same biological resource (same cell type or tissue) can be used to accomplish the back-phosphorylation (Li et al., 2016; Mundina-Weilenmann et al., 1991) without any pre-dephosphorylation process(Tsai et al., 2015). *In vitro* kinase reactions both at the protein and tryptic peptide levels can be used in most cases, except for basophilic kinase reactions, where tryptic peptides cannot be used as substrates due to the lack of K or R at the N-terminal side of the phospho accepting sites. In such a case, the kinase reaction at the protein level, followed by tryptic digestion, is used to generate the basophilic motif-centric phosphopeptides. After TMT labeling, phosphopeptides are enriched by metal affinity chromatography and analyzed by nanoLC/MS/MS (Figure 1b). The TMT-labeled precursor ions from the endogenous and back-phosphorylated peptides are fragmented and the assembled signals in b or y ions are helpful for peptide identification, especially for endogenous phosphopeptides with low abundance. The relative quantitation of endogenous phosphopeptides across different samples is done by the use of reporter ions at the MS^3^ level, by the implementation of synchronous precursor selection (SPS)-based MS^3^ technology (McAlister et al., 2014), which can reduce the interference signals due to co-isolation of precursor ions.

**Figure 1.**
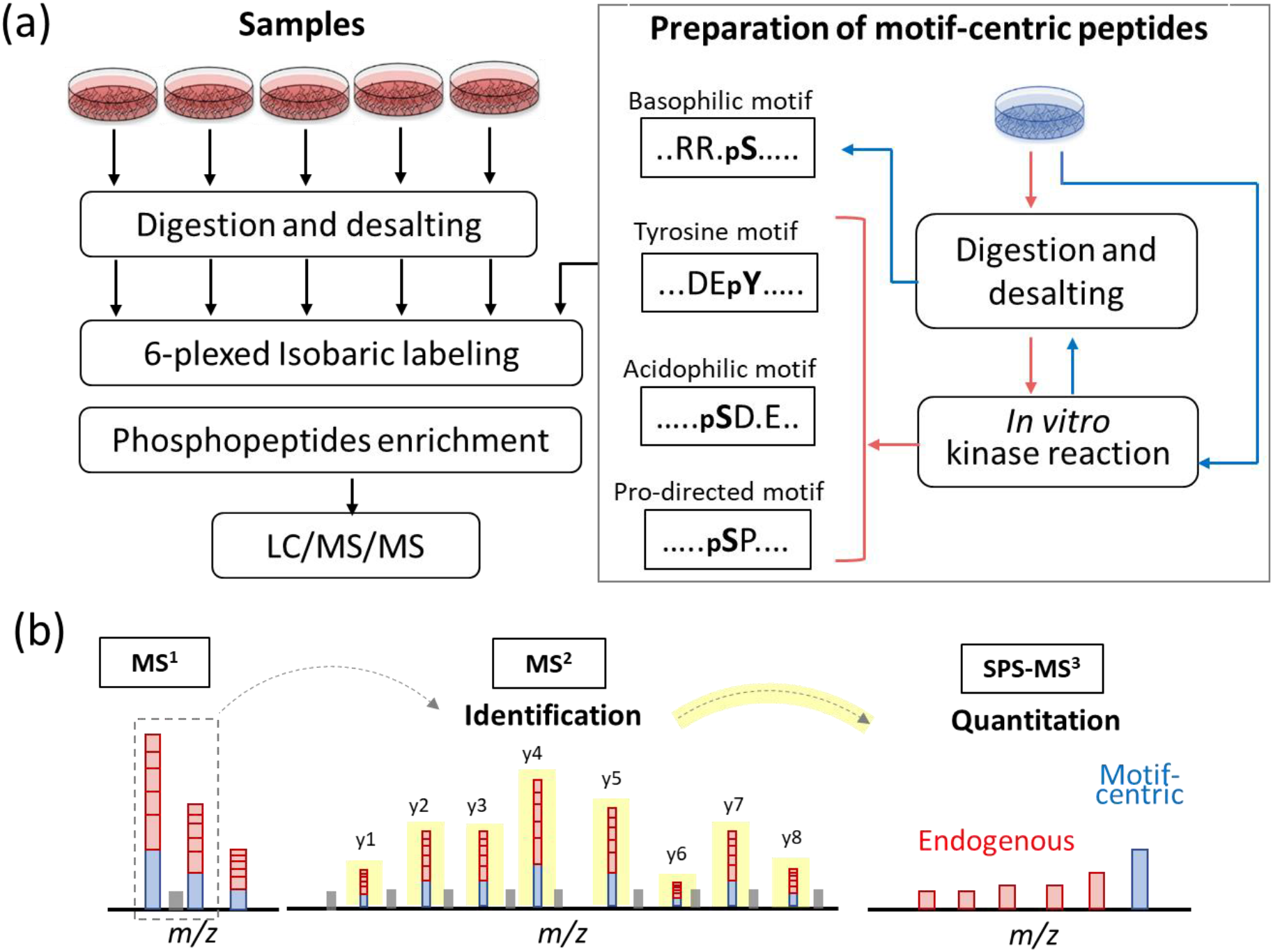
Workflow of isobaric motif-centric phosphoproteomics with *in vitro* kinase reaction. (a) The motif-centric peptides were generated by *in vitro* kinase reaction. Tryptic peptides from study samples and motif-centric peptides were labeled with different isobaric tags (TMT in this study). After mixing, the TMT-labeled phosphopeptides were enriched and analyzed by LC/MS/MS. (b) TMT-labeled ions from endogenous and motif-centric phosphopeptides are assembled as a single peak at the MS^1^ level and then fragmented at the MS^2^ level to identify the peptide sequence. Relative quantification of endogenous phosphopeptides between different samples is achieved by reporter ions at the MS^3^ level.

### Acidophilic motif-centric phosphoproteome analysis

We firstly employed the motif-centric TMT approach to quantify the kinase-perturbed phosphorylation changes in HeLa cells using CK2 kinase inhibitor (CKi, Silmitasertib, CX-4945). The tryptic peptides from CKi-treated HeLa cells were phosphorylated by CK2 *in vitro* and labeled with both TMT^128^ and TMT^131^ for CK2 motif-centric boosting channels. The phosphopeptides from DMSO-treated cells were labeled with both TMT^126^ and TMT^129^, and phosphopeptides from CKi-treated cells were labeled with both TMT^127^ and TMT^130^. An MS^2^ spectrum and an MS^3^ spectrum of a known CK2 substrate, the pS66 site in LIG1, are shown in Figure 2. After fragmentation by CID, the peptide sequence and phosphosite localization information can be annotated at the MS^2^ level (Figure 2a). Then, the MS^3^ spectrum was obtained, demonstrating that the TMT signals of endogenous phosphopeptides were decreased after the CKi treatment (TMT^127^ and TMT^130^ in Figure 2b) and that the TMT signals of CK2-motif targeting phosphopeptides were increased after *in vitro* kinase reaction (TMT^128^ and TMT^131^ in Figure 2b). These results indicate that this isobaric motif-centric approach can be used to monitor CK2 phosphorylation signaling in terms of the TMT ratios.

**Figure 2.**
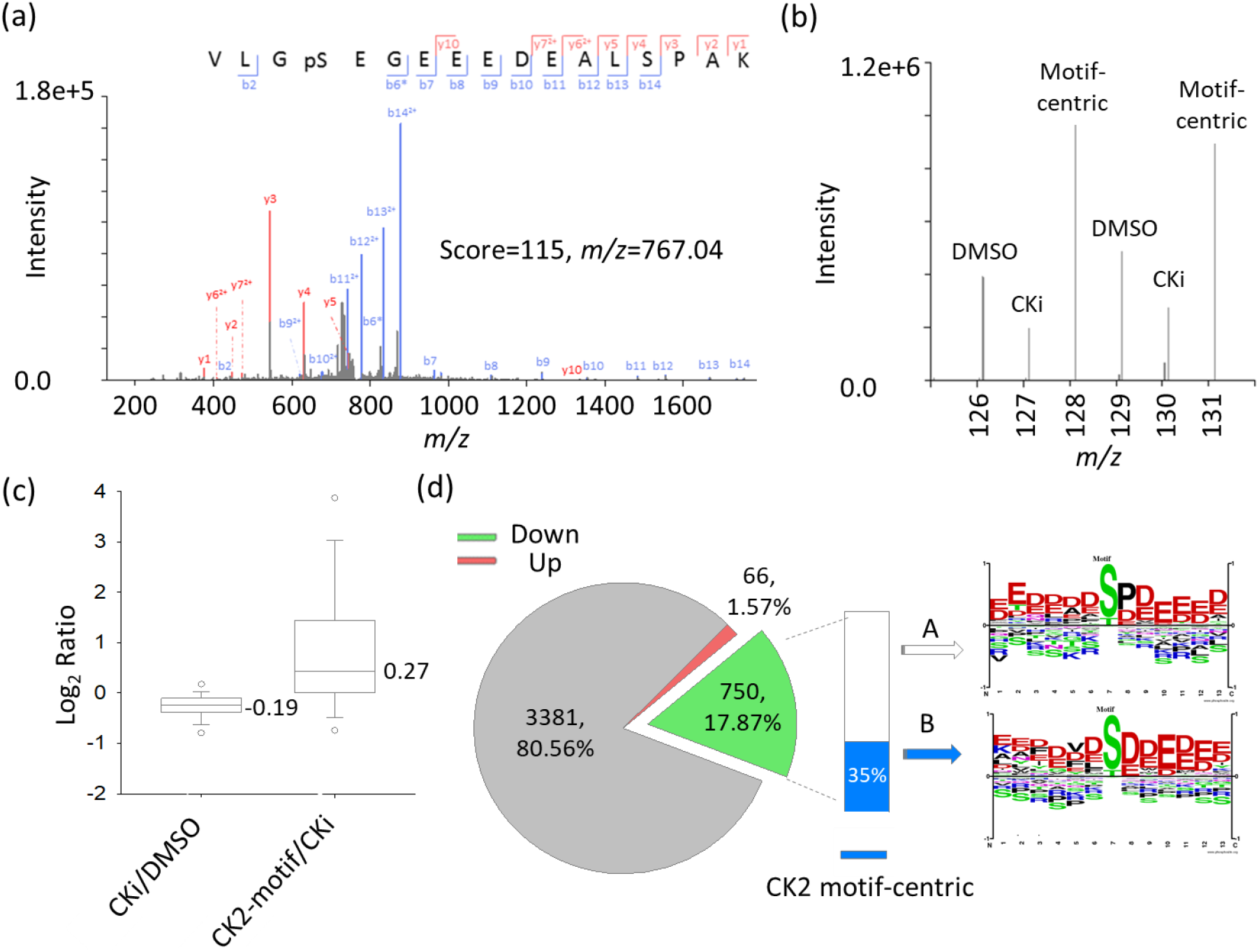
Typical example of the isobaric CK2 motif-centric phosphoproteomic approach. (a) An MS/MS spectrum of TMT-labeled phosphopeptide (VLGpSEGEEEDEALSPAK, triply charged). Product ions at the MS^2^ level identified the peptide sequence and phosphorylation site localization. (b) An MS/MS/MS spectrum at the 6-plexed TMT reporter ion region. Three samples (DMSO, CKi-treated and CK2 motif-centric phosphopeptides) in duplicate preparations, labeled with 6-plexed TMT reagents, were quantified at the MS^3^ level. (c) The ratio distribution on a log_2_ scale of identified phosphopeptides. Left bar: CKi/DMSO, right bar: motif-centric/CKi. (d) Quantitation result for CKi treatment. Sequence motif analysis for the down-regulated phosphorylation sites after CKi treatment was performed. The blue bar indicates the amount of phosphorylation sites significantly increased after in vitro kinase reaction (CK2-motif centric peptides). (e) The log_2_ ratio correlation between CKi/DMSO and motif-centric/CKi. The red and green circles mean up-regulated and down-regulated phosphorylation sites after CKi treatment, respectively, in panel (d). The blue fill in the green circle indicates downregulated CK2-motif centric peptides in panel (d).

In total, this isobaric motif-centric approach quantified 4559 unique phosphopeptides (91% specificity in phosphopeptide enrichment) from 25 μg of starting material. The logarithm of the median ratio of CKi-treated to untreated peptides was negative, while the logarithm of the median ratio of motif-centric (CK2-motif) to CKi-treated peptides was positive and this increase was larger than the decrease caused by CKi treatment (Figure 2c), indicating that the *in vitro* kinase reaction effectively increased the TMT signals of peptides directly phosphorylated by CK2. Silmitasertib has been used to exclusively inhibit CK2 activity in previous studies(Chon et al., 2015; Wang et al., 2017). Therefore, we can discriminate direct CK2 substrates from others by comparison of TMT ratios such as CKi/DMSO and CK2-motif/CKi using Student’s t-test. Based on the CKi/DMSO ratios, 66 (1.6%) up-regulated and 750 (18%) down-regulated phosphorylation sites were identified (Figure 2d). Among the down- regulated phosphorylation sites, up to 35% (n = 266) were significantly increased after *in vitro* kinase reaction (group B in Figure 2d). After sequence motif analysis (O’Shea et al., 2013), we found that the sequence motif of the phosphopeptides in group B agreed with the known CK2 substrate motif (acidic motif), whereas the motif logo from group A contained both Pro and Asp at +1 position (Figure 2d). Among the phosphorylation sites (n = 266) in group B, 222 sites were registered in a public phosphorylation sites database (Hornbeck et al., 2015), including 6 known CK2 substrates: PTGES3 (S113), LIG1 (S66), SLC3A2 (S375), CDK1 (S39), TOP2A (S1377) and ABCF1(S110). For group B proteins, we also examined the overlap with CK2 interactors in STRING and found 20 known CK2 interactors (Figure S1a). We further performed Gene Ontology and Reactome Pathway enrichment analysis for group B proteins(Szklarczyk et al., 2019). As a result, 137 proteins in group B were annotated as nuclear proteins, and a majority of them possessed functions related to ATP binding, DNA binding, nucleotide binding and so on (Table S1). Among the annotated pathways, the top hit was the cell cycle pathway, in which 21 group B proteins were down-regulated, including CDK1, a known CK2 substrate that affects cell cycle regulation upon S39 phosphorylation(Bloom and Cross, 2007; Russo et al., 1992) (Figure S1b). All these results indicate that this motif-centric approach is an effective tool for monitoring specific kinase-mediated signaling pathways.

### Basophilic motif-centric phosphoproteome analysis

The isobaric motif-centric approach was further examined with PKA as a basophilic kinase, using forskolin as an activator. Unlike acidophilic CK2, PKA cannot be used to generate motif-centric peptides from tryptic peptides because it requires basic amino acid residues at the N-terminal side of the phospho-acceptor. Therefore, PKA motif-centric peptides were generated by *in vitro* kinase reaction at the protein level, followed by tryptic digestion (Fig. 1a). After the isobaric labeling, the TMT ratios of three channels, forskolin-treated, DMSO-treated and motif-centric samples, were used to discriminate the peptides phosphorylated directly by PKA from others, as in the case of CK2. The results are illustrated in Figure S2. In total, we quantified 7855 phosphorylation sites from biological replicate experiments. Among them, 620 phosphorylation sites were upregulated (group 1) and 322 phosphorylation sites were inhibited by forskolin (group 2). We previously established a computational model (primary sequence preference, PSP score) to characterize the kinase sequence specificity toward the substrate target site based on known kinase-substrate relationships, in order to exclude potential indirect targets of PKA (Imamura et al., 2017). We applied PSP scoring to the phosphorylation sites in groups 1 and 2, and found that the PSP scores of group 1 were significantly higher than those of group 2, although the sequence motifs of phosphorylation sites in groups 1 and 2 both belonged to the basophilic category (Figure S2b). Based on the above results, the motif-centric approach was able to discriminate peptides phosphorylated directly by PKA from indirectly phosphorylated ones.

### Tyrosine motif-centric phosphoproteome analysis

Although the detectability of pY sites has been restricted by their extremely low phosphorylation stoichiometry compared to pS and pT sites(Sharma et al., 2014; Tsai et al., 2015), this lower stoichiometry results in a larger amount of unphosphorylated counterparts, which can be used for back-phosphorylation. In order to increase the identification number of pY sites, we firstly tried to use pervanadate (PV)-treated Hela cells as a pY-centric sample (Figure S3). PV is well-known as a tyrosine phosphatase inhibitor causing an increase in the stoichiometry of endogenous pY sites(Sharma et al., 2014). As expected, the pY content was increased from 0.8% to 11.7% (Figure S4a) by PV treatment. Then we employed the PV-treated peptides as pY-centric peptides with 25 μg of untreated HeLa peptides. Indeed, the number of quantifiable pY sites increased from 74 to 595 as the spiking amount of the pY- centric peptides was increased from 0 to 75 μg, without any antibody-based enrichment (Figure S4b). The amplified signals caused by the spiked pY peptides resulted in a much larger number of identified pY sites than in usual phosphoproteome analysis. However, the number of identified pY sites was still lower than that obtained using the recently published antibody-based boosting strategy (Chua et al., 2020), which also used PV-treated cells as boosting samples to quantify over 2300 unique pY peptides from 1 mg of starting materials.

In order to extend the tyrosine phosphoproteome coverage, the back-phosphorylation sites of the untreated HeLa peptides were phosphorylated via *in vitro* kinase reaction by using tyrosine kinases such as EGFR and SRC. Note that EGFR and SRC have different phosphorylation motifs (Sugiyama et al., 2019). Then, the kinase-treated peptides were labeled with one of the 6-plexed TMT reagents to detect the endogenous pY sites in EGF or EGF/afatinib (EGFR inhibitor)-treated Hela cells (Figure S3). After IMAC enrichment, the TMT-labeled phosphopeptides were analyzed on a 2-m long monolithic silica column system with the SPS-MS^3^ technique (McAlister et al., 2014). Compared with the result from PV-treated HeLa as pY motif-centric peptides, the content of pY increased from 15% to 87% (Figure 3a) which resulted in an approximately 10-fold increase in the numbers of identified and quantified Class 1 pY sites (localization probability >0.75), as shown in Figure 3b. However, Cheung *et al*. (Cheung et al., 2021) reported that the isobaric labeling-based quantitative approaches have technical limitations that potentially affect data quality and biological interpretation, due to the large amounts of spiked carrier samples. In addition, it is difficult to control the TMT ratio within the quantifiable range because the phosphorylation stoichiometry in cells depends on each pY site. Furthermore, we should reject TMT-peptides without reporter ion signals in the sample channels. Therefore, we examined the distribution of the TMT reporter ion intensity of each sample channel and found a notch to discriminate the signal from the noise (Figure S5); this was also mentioned in the previous study (Hughes et al., 2017). Based on this observation, we set the acceptance criterion for the minimum TMT intensity in the sample channels to be greater than 40 on a log_2_ scale for the total TMT intensity of the sample channels. We also set another criterion, that the maximum TMT ratio of the motif-centric channel to the sample channel should be less than 100 (Figure 3c). By applying these two criteria, we found that 668, 5425 and 3688 pY sites could be quantified in the PV-treated, EGFR-centric and SRC-centric samples, respectively. Among these pY sites, the sequence motifs of unique pY sites in each dataset were different (Figure 3d), which indicated that complementary pY sites can be identified by using different motif-centric peptides. For quantitation based on the TMT reporter ion intensities, the EGF and EGF-TKI treated cells were separated in the PCA analysis (Figure 3e). The above results demonstrate that this motif-centric approach through spiking isobaric tyrosine phosphopeptides with high purity was able to increase the detectability of pY sites without the need for any affinity purification procedure, such as application of pY antibody.

**Figure 3.**
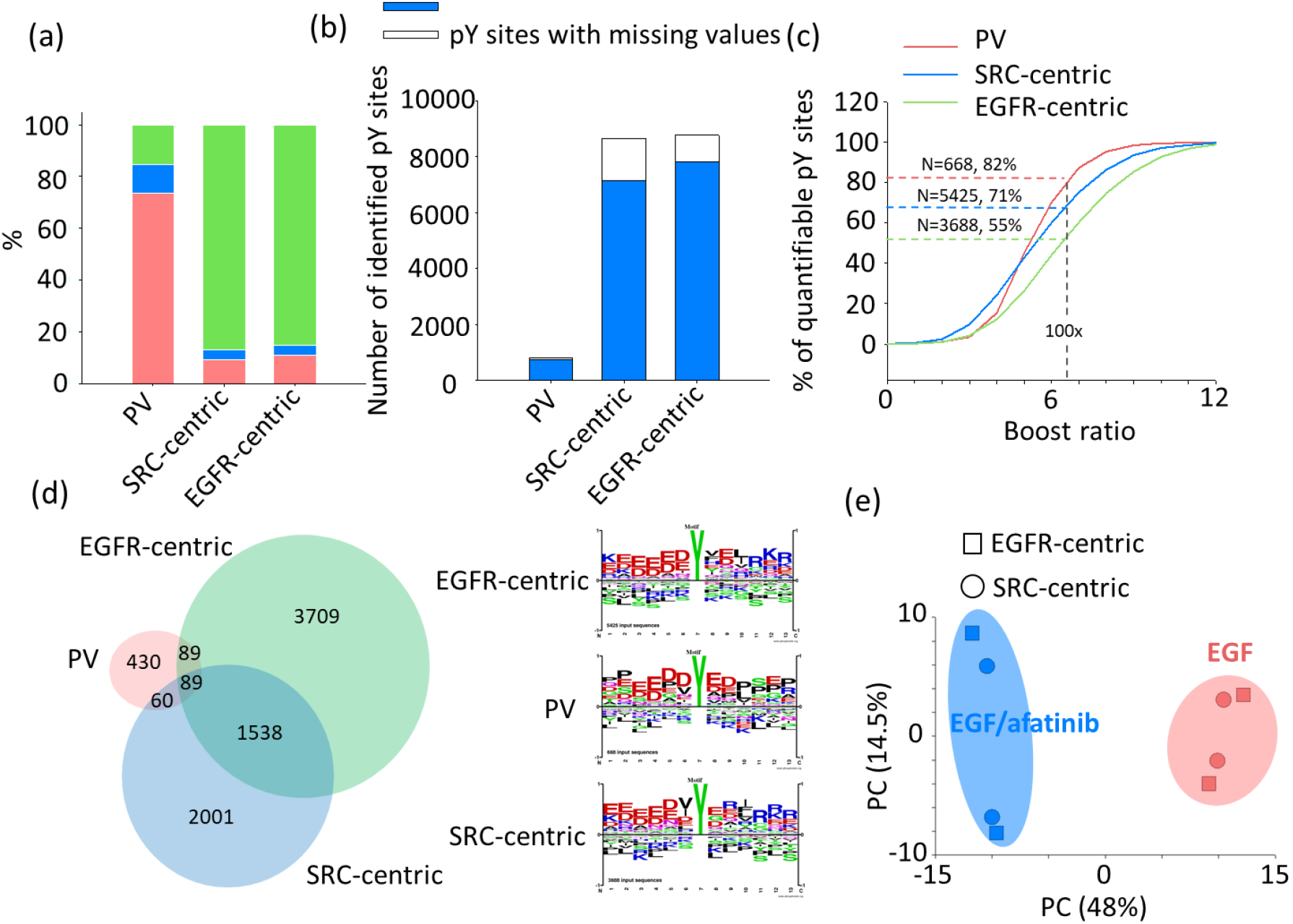
Summary of pY motif-centric phosphoproteomic approach. (a) The content (%) of identified pY sites using PV-treated, SRC and EGFR phosphorylated peptides as the pY-centric samples. (b) The number of pY sites identified in each experiment. A quantifiable site is defined as a pY site having signal intensities in at least 2 TMT reporter channels. Otherwise, pY sites were considered as missing values. (c) The content of quantifiable pY sites filtered by boosting ratio, defined as the TMT signal at *m/z* 131 divided by the averaged signal at *m/z* 127 and 129. (d) The overlap and the sequence logos of boosting ratio-filtered quantified pY sites between different motif-centric approaches (e) The PCA analysis of commonly quantified pY sites (filtered).

### Multiple motifs-centric approach to depict perturbed phosphoproteome

We further used multiple kinases to generate a wide variety of motif-centric peptides to monitor the phosphorylation signals generated by different kinases. For proof-of-concept, we analyzed the phosphoproteome of EGF-treated and EGF/afatinib co-treated HeLa cells. We selected four Pro-directed kinases ((ERK1 (MAPK3), JNK1(MAPK8), p38α (MAPK14) and CDK1)), two tyrosine kinases (SRC and EGFR) and one acidophilic kinase (CK2) to detect the different motif-centric phosphorylation sites (Figure S6). After filtering based on the boosting ratio (<100x) and TMT intensity cut-off (sum of TMT intensity from sample channels >40 on log2 scale), up to 11,895 class 1 phosphorylation sites were quantified (at least two valid TMT values in one cell type) including 5,045 pS, 1,756 pT and 5,094 pY sites (Figure 4a). According to Student’s t-test (EGF vs. EGF/afatinib), the ratio of commonly regulated phosphorylation sites was consistent between experiments using different kinases, regardless of the kinase used (Figure 4b), indicating that the boosting channel did not significantly affect the quantitative results.

**Figure 4.**
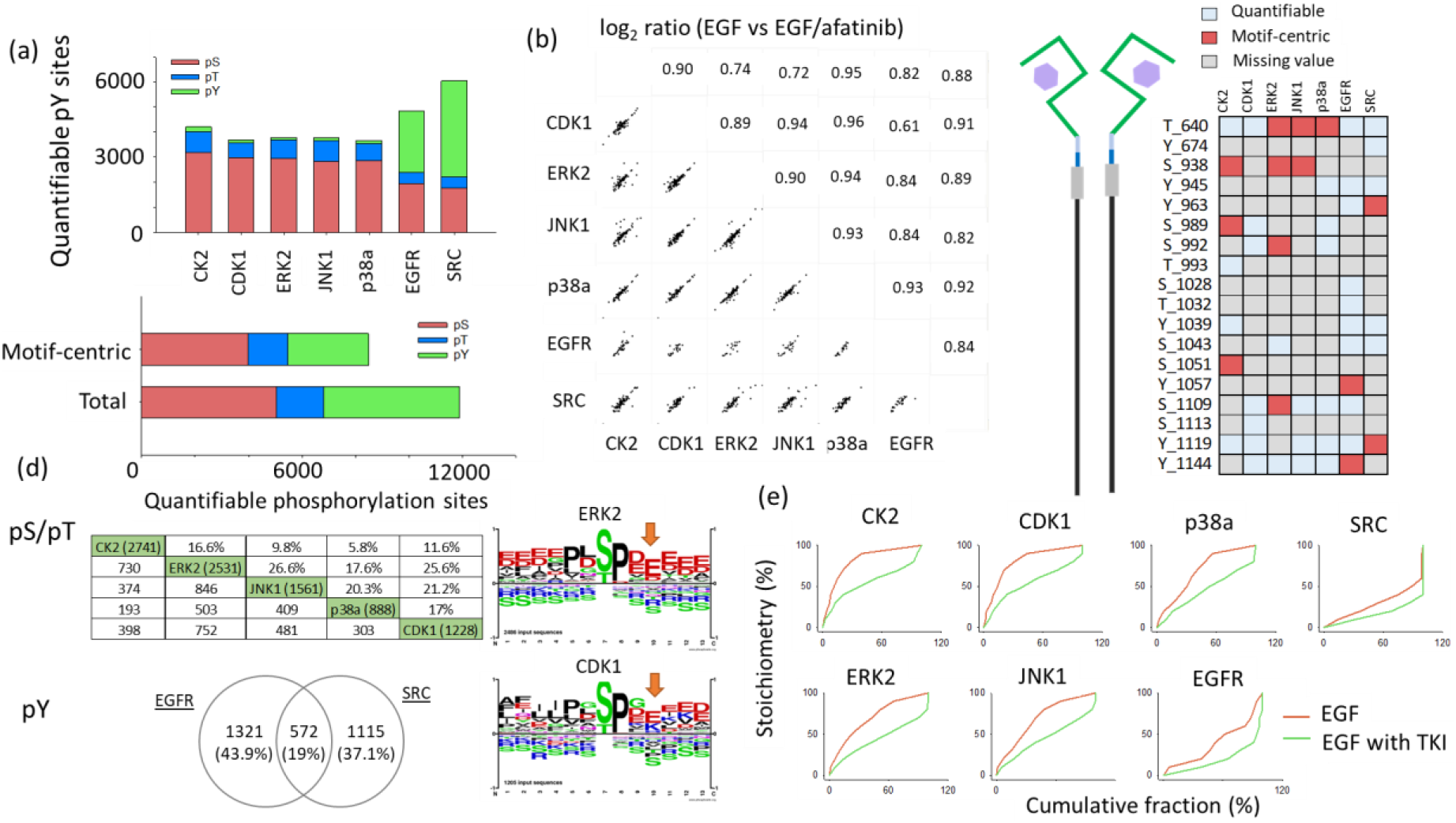
Summary of motif-centric phosphoproteomic approach with multiple kinases. (a) The number of quantified phosphorylation sites (class 1) obtained using different kinases. (b) The ratio correlation of commonly regulated phosphorylation sites (EGF vs. EGF/afatinib, p<0.05) in experiments with different kinases. (c) The motif-centric sites (red color) among quantified phosphorylation sites in EGFR. (d) The overlap of motif-centric phosphorylation sites between the samples with different kinases as motif-centric samples. The sequence motif analysis for the ERK2 and CDK1 centric phosphorylation sites. (e) The phosphorylation stoichiometry distribution of motif-centric phosphorylation sites after afatinib treatment.

In addition to identifying which phosphorylation sites were regulated after afatinib treatment, we were also able to estimate which kinases phosphorylated which sites by using this motif-centric approach. For example, up to 18 class 1 phosphorylation sites, including 7 pY sites, on EGFR kinase were quantified without any immunoprecipitation step from 25 μg of starting materials per TMT channel (Figure 4c). Among these sites, pY1144 is a known autophosphorylation site which was downregulated by afatinib and detected in the EGFR motif-centric experiment. This result indicates that the activity of this phosphorylation site is controlled by EGFR and also inhibited by afatinib. On the other hand, pS1051 and pS992 were downregulated by afatinib, but there is no information about the corresponding kinase in the public database. Through this motif-centric approach, we could determine that pS1051 and pS992 are likely phosphorylated by CK2 and ERK2, respectively.

Another advantage of the motif-centric approach is the specificity of the *in vitro* kinase reaction. Student’s t-test (EGF/afatinib vs. motif-centric) revealed a low overlap of motif-centric peptides among the kinases (Figure 4d). Although ERK2, JNK1, p38a and CDK1 are all Pro-directed kinases, complementary profiles for the quantified motif-centric phosphorylation sites were observed (Figure 4d). The motif logos of these sites showed slight differences among the four Pro-directed kinases (Figure S7a). For example, the proportion of pT motifs was higher with p38α kinase. In addition, more acidic amino acids were located at the C-terminal side of the phospho-acceptor site in the case of ERK2 kinase compared to CDK1 kinase. In addition to S/T sites, the pY motifs also differed between EGFR- and SRC-centric phosphopeptides (Figure S7b).

Further, from the two ratios (EGF vs. motif-centric and EGF/afatinib vs. motif-centric), we can estimate the stoichiometry of motif-centric phosphorylation sites (Figure 4e), assuming that the efficiency of *in vitro* kinase reaction is 100%. The phosphorylation sites containing acidophilic kinase substrates targeted by CK2 generally exhibited higher phosphorylation stoichiometry than sites targeted by Pro-directed kinase and tyrosine kinases. The phosphorylation stoichiometry distribution that we observed here is consistent with our previous findings(Tsai et al., 2015). To further validate that the endogenous signals boosted by motif-centric peptides are lower than other peptide signals, the peak areas in the XICs of MS1 signals of quantified phosphopeptides were calculated based on the proportion of TMT intensity (Figure S8). The XICs of endogenous phosphopeptides boosted by the motif-centric peptides were lower than those of other phosphopeptides, which indicates the motif-centric approach is effective to identify these low-abundance kinase substrates.

In this motif-centered approach, a specific kinase can be chosen to target an endogenous peptide that is phosphorylated by that kinase. To confirm this, 8482 motif-centric phosphopeptides prepared with these seven kinases were subjected to KEGG pathway enrichment analysis using DAVID(Huang da et al., 2009). The results showed that, as expected, known pathways involving the seven kinases, such as ErbB and insulin signaling, were enriched (Figure 5a and Table S2). We then performed STRING protein-protein interaction analysis of the 33 phosphoproteins comprising the ErbB pathway identified in this enrichment analysis (Figure 5b). Furthermore, we mapped the responsible kinases for the *in vitro* phosphorylation sites on these proteins (Figure 5c), showing how the kinases used for motif-centric peptides covered the targeted phosphosites within the targeted pathway. In other words, we can manipulate the targeted pathway by choosing the appropriate kinases, using our large-scale library of *in vitro* kinase-substrate relationships(Sugiyama et al., 2019). Overall, our findings indicate that the motif-centric approach can provide system-wide customizable maps consisting of targeted pathways under physiological or pathological regulation.

**Figure 5.**
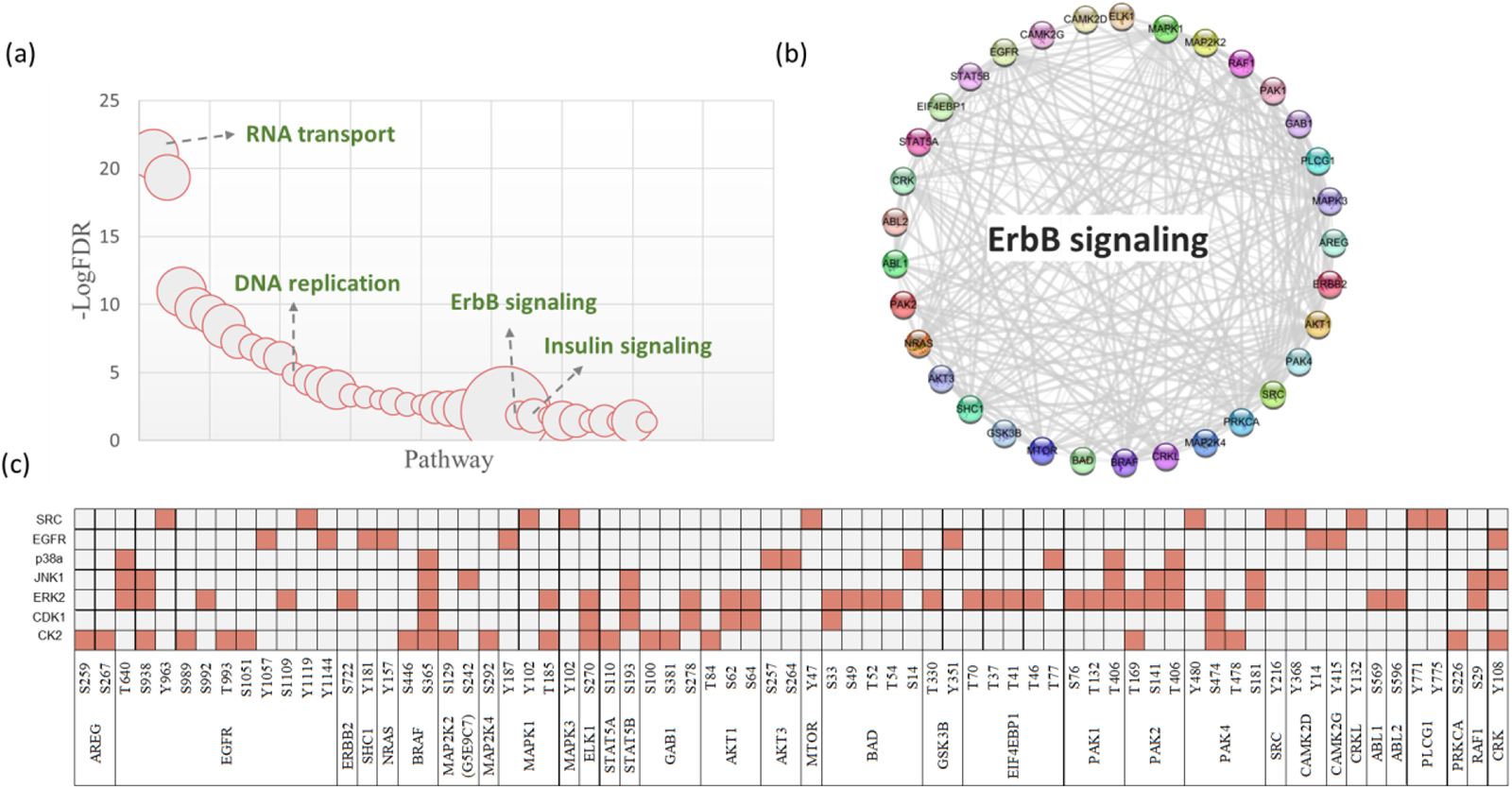
Pathway enrichment analysis of motif-centric phosphorylated sites with multiple kinases. (a) KEGG pathway enrichment analysis by DAVID. (b) STRING protein-protein interaction analysis of the phosphoproteins enriched in the ErbB signaling pathway. (c) Phosphorylation sites on proteins in the ErbB signaling pathway and the corresponding kinases.

## Discussion

The kinase assay-linked phosphoproteomics (KALIP) approach(Xue et al., 2013; Xue et al., 2012) has been developed to find potential kinase substrates. Bona fide direct substrates were found in the overlap of the *in vitro* phosphopeptides generated by kinase reaction with dephosphorylated peptides and *in vivo* kinase-dependent phosphorylation events in different LC-MS/MS runs. However, the dephosphorylation is not complete, and this may result in the false-positive identification of putative kinase substrates. Although stable-isotope KALIP(Xue et al., 2014) using ^18^O-ATP as a phosphate donor instead of ^16^O-ATP for *in vitro* kinase reaction improved the situation, the analytical throughput is still limited by the need for multiple LC/MS/MS analysis, and in addition the background effect due to incompletely dephosphorylated peptides reduces the detection sensitivity for putative kinase substrates. To tackle these challenges, we previously established a computational model (primary sequence preference, PSP score) to characterize the kinase sequence specificity toward substrate target sites based on known kinase-substrate relationships in order to exclude potential indirect targets of PKA (Imamura et al., 2017). Furthermore, the isobaric motif-centric approach developed in this study can link *in vitro* substrates and physiological phosphorylation events by monitoring both endogenous and back (specific kinase motif-targeting) phosphorylated signals in a single LC/MS/MS run without the need for a further dephosphorylation step, thereby enabling high-throughput analysis of putative kinase substrates. For example, the isobaric motif-centric approach was able to detect 266 CK2 motif-centric phosphorylation sites downregulated by CKi (Figure 2d). This method not only provides information about the fold-change between different samples, but also allows estimation of the stoichiometry of motif-centric phosphorylation sites (Figure 4e) based on the ratio between endogenous and motif-centric back phosphorylated signals, assuming that the efficiency of kinase reaction is 100%.

Unlike metal affinity chromatography for enrichment of phosphopeptides, motif-specific immunoaffinity precipitation (IAP)-based LC/MS/MS makes it possible to recognize a characteristic sequence motif from a broad range of peptides by using different motif antibodies. Because each antibody binds phosphopeptides followed by a specific peptide motif, the overlap of identified phosphopeptides among different antibodies is low. The results of this approach are similar to those obtained with our motif targeting approach (Figure 4d), in which recognition between kinase and peptides is based on the specific sequence motif(Stokes et al., 2015; Sugiyama et al., 2019). The overlap between a given antibody and metal affinity chromatography ranged from roughly 16% with the all Ser/Thr antibody mix, to a low of only 3.6% with phosphotyrosine pY-1000 (Stokes et al., 2015). Anthony *et al*. aimed to enlarge the phosphoproteome coverage by using both TiO_2_ followed by basic pH reversed-phase fractionation and motif-specific IAP with four different phosphorylation motif-specific antibodies(Possemato et al., 2017). In total, 8947 nonredundant peptides were identified in the TiO_2_ data set, of which only 852 (9.5%) were in common with the peptides identified in the IAPs. These results suggest that the range of phosphorylation within a given system is so broad that no single approach is likely to provide comprehensive coverage.

In the case of IAPs, the specificity of the antibodies is not high enough to distinguish phosphopeptides with similar motifs; for example, MAPK phosphorylates substrates with the consensus sequence PX(S/T)P and CDKs phosphorylate substrates containing the consensus sequence (S/T)PXR/K(Songyang et al., 1996). However, the complementary regulated motif-centric phosphorylation sites (Figure 4d and Figure S7) were differentiated in this study, even though the kinases all belong to the same Pro-directed kinase group. The motif-centric approach was also able to distinguish the motif difference (PX(S/T)P and (S/T)PXR/K) for ERK2 and CDK1 (Figure 4d). In addition, the purification specificity in IAP (less than 50%) is much lower than that in the metal affinity-based approach (Possemato et al., 2017), which means that mg amounts of starting materials are necessary for the IAPs method. For our isobaric motif-centric approach, we performed metal affinity chromatography (IMAC in this study) after motif-centric enrichment, and obtained a phosphopeptides purification specificity of over 90% (Figure S9). In addition, the IAPs approach uses selected antibodies to isolate targeting peptides, and this may cause unnecessary sample loss during the purification step. In contrast, the motif-centric approach uses kinase to recognize specific substrates and transfer the phosphate group to back-phosphorylated peptides. Endogenous phosphopeptides, especially low-abundance tyrosine phosphopeptides, are not removed before IMAC enrichment. Therefore, only a few tens of μg of material was required for our isobaric motif-centric approach. Recently, a novel antibody-based method, called PTMScan Direct, was developed for the identification and quantitation of peptides derived from proteins that are critical signaling nodes of various pathways (Stokes et al., 2012). However, the coverage of the IAPs approach is still limited by the availability and quality of antibodies. In contrast, recombinant active kinases are much more readily available than antibodies. In our previous study (Sugiyama et al., 2019), we were able to identify a total of 175,574 potential direct kinase substrates by using 385 active kinases (354 wild-type protein kinases, 21 mutants and 10 lipid kinases). Based on this kinase substrate library, it is easily possible to select multiple kinases for targeting pathway analysis.

The multiplexing nature of isobaric labeling is particularly useful to achieve higher sensitivity with limited individual sample amounts, as in the case of tyrosine phosphopeptides. Chua *et al*. (Chua et al., 2020) and Fang *et al*. (Fang et al., 2020) used samples with and without pervanadate (tyrosine phosphatase inhibitor)-treated cells as a boosting channel to increase the relative abundance of tyrosine phosphorylation sites. Although around 2300 (Chua et al., 2020) and 835 pY phosphopeptides (Fang et al., 2020) were detected, the required amount of starting material is at the mg level. In addition, the usage of antibodies for pY phosphopeptides enrichment is also necessary for their approach. Here, in contrast, we generate higher-purity tyrosine phosphopeptides via *in vitro* kinase reaction (motif-centric). By using motif-centric peptides, we could detect up to 7129 (SRC targeting) and 7280 (EGFR targeting) tyrosine phosphopeptides without the need for immunoprecipitation, using only a few tens of μg of starting materials.

## Limitations

Although the performance of isobaric labeling-based quantitative approaches is affected by co-selected precursor ions, the usage of SPS-MS^3^ can minimize this effect and improve the detection sensitivity by utilizing co-fragmented multiple (up to 10) MS^2^ fragment ions with higher intensity (McAlister et al., 2014). Moreover, the newly available Real Time Search-MS^3^ method (RTS-MS^3^) (Erickson et al., 2019) or ion mobility technique (Bekker-Jensen et al., 2020; Hebert et al., 2018; Ogata and Ishihama, 2020) provides a solution for precise and accurate quantitation without sacrificing proteome coverage. Recently, the use of spiked TMT labeling peptides with boosting or carrier samples has reduced the TMT reporter ions dynamic range and decreased the quantitation accuracy (Tsai et al., 2020). Cheung et al. (Cheung et al., 2021) and Stopfer et al. (Stopfer et al., 2021a) also demonstrated that an increase in carrier proteome level requires a concomitant increase in the number of ions sampled to maintain quantitative accuracy. Therefore, it will be important to optimize the spiking amount of carrier/boosting and the TMT channel design for the isobaric motif-centric strategy. Optimization of the MS parameters to improve the ion sampling by adjusting the ion injection time and AGC will also be beneficial to improve the quantitation performance(Cheung et al., 2021; Tsai et al., 2020). To overcome the challenge presented by the large quantitation dynamic range, it may be useful to integrate the motif-centric approach with isotope-based SureQuant (Stopfer et al., 2021b) or Internal Standard Triggered-Parallel Reaction Monitoring (IS-PRM) (Gallien et al., 2015) quantitation.

## Conclusions

The isobaric motif-centric strategy presented here can be used to enhance the sensitivity of specific kinase downstream signaling analysis, especially for tyrosine phosphopeptides. It provides a simple yet highly effective quantitative phosphoproteomic workflow suitable for multiplexed analysis of relatively small biological or clinical samples (sub mg), including cells or tissues. This approach enables the quantitation of both fold-change and stoichiometry among thousands of phosphopeptides generated by specific kinases in signaling pathways. The usage of multiple kinases for motif targeting analysis effectively increases the phosphoproteome coverage. Overall, we anticipate this strategy should find broad biomedical applications for targeting kinase/pathway analysis where limited amounts of starting cells or tissues are available.

## Acknowledgement

We would like to thank all lab members for fruitful discussions. C.-F.T. was supported by a JSPS Grant-in-Aid for a postdoctoral fellowship for overseas researchers (15F15343). This work was supported by the JST Strategic Basic Research Program, CREST (grant No. 18070870) to Y. I., and by a JSPS Grants-in-Aid for Scientific Research (No. 17H03605, No. 21H02459 to Y. I., No. 20K21478 to Y. I. and K. O., and No. 20H04845, No. 21H02466 to N. S.)

## Author Contributions

Conceptualization, C.F.T. and Y.I.; Methodology, C.F.T. and Y.I.; Investigation, C.F.T., K.O. and N. S.; Writing – Original Draft, C.F.T.; Writing –Review & Editing, C.F.T. and Y.I; Funding Acquisition, Y.I.; Supervision, Y.I.

